# A zebrafish knock-in reporter line for the Foxo1a transcription factor

**DOI:** 10.1101/2023.07.17.548093

**Authors:** Inés Garteizgogeascoa Suñer, Sumeet Pal Singh

## Abstract

Zebrafish is a powerful model organism for in vivo live imaging. However, protein visualization relies to this date on the overexpression of fluorescently tagged proteins from a specific promoter which very often does not recapitulate endogenous patterns of expression and dynamics. Tagging proteins in the endogenous locus is not widely used in the field due to its technical inefficiency and difficulty. Here we developed a knock-in reporter line for the Foxo1a transcription factor by inserting an EGFP-pA cassette in frame at the C-terminal generating a fusion protein. Foxo1a has been involved in the regulation of many biological processes regarding metabolism, stress response, longevity, cell differentiation and others and its functions are conserved from invertebrates to vertebrates. Using in-vivo live imaging at early developmental stages, we validated the expression in the cardiovascular network, CNS, olfactory epithelium, spinal cord, retina, skeletal muscle, and myocardium. Our line opens the way for imaging studies aiming to characterize the expression and localization of this transcription factor in a tissue and context specific manner. Also, the knock-in line can be used in combination with other modern techniques to determine the transcriptional targets of Foxo1a, many of which remain unknown.

## MAIN TEXT

The forkhead box protein O1 (FOXO1) is one of the four members of the FOXO subfamily of forkhead transcription factors. FOXO proteins are master signaling integrators that modify gene expression upon a big diversity of environmental stimuli. For this, they have been involved in the regulation of many biological processes such as development, longevity, metabolism, apoptosis and stress response (Alikhani et al., 2007; Greer & Brunet, 2005; Rached et al., 2010; van der Vos & Coffer, 2011). The different environmental stimuli are coded into the “FOXO code” a combination of post-translational modifications (PTMs) which can include phosphorylation, methylation, acetylation and ubiquitylation and determine the nuclear/cytoplasmic trafficking of FOXO proteins (Y. Wang et al., 2014). Once in the nucleus, they transcriptionally control a wide range of downstream targets in peripheral tissues but also in the central nervous system (CNS) (Gilels et al., 2013; Hoekman et al., 2006; Polter et al., 2009). For example, one of the main regulatory mechanisms described for FOXO factors is their regulation by the PKB/Akt pathway. PKB/Akt inhibits the transcriptional activity of FOXO members by causing their redistribution from the nucleus to the cytoplasm in response to phosphorylation (Kops et al., 1999). Even though on many occasions all four members are treated as a family, each of the FOXO proteins can be expressed in a spatially and temporally restricted manner (Hoekman et al., 2006; Jacobs et al., 2003) In zebrafish, there are two orthologs of the mammalian *Foxo1* gene, termed *foxo1a* and *foxo1b* which share 53 and 51% identity with human *FOXO1*, respectively.

The zebrafish is an important vertebrate model organism. However, research using this model has been historically influenced by the lack of good antibodies that work either on wholemount or fixed tissue. This is primarily because most of the zebrafish community relies on the cross-reactivity of mammalian antibodies which is typically impaired by the lack of antigen conservation and of protein specificity. Despite a modest growing number of specifically raised antibodies, there is a need to develop new tools that allow the visualization of proteins in zebrafish.

The solution to this limitation has been and continues to be the generation of overexpression systems by promoter-driven transgenesis where either the whole or a fragment of the coding sequence of a desired gene is linked to a fluorescent protein (FP) and subcloned to a plasmid that contains either the specific promoter of the gene, if it is known or a strong specific promoter that targets the expression specifically to a cell type or tissue of interest This construct is then used to generate a stable transgenic line by injection at a single-cell stage. This notion relies on the fact that the expression of such desired gene is detected at mRNA level by in situ hybridization or transcriptomic techniques like bulk or single-cell RNAseq although studies have shown spectacularly poor correlation between mRNA and protein expression levels, creating concern for inferences from only mRNA expression data (Abreu et al., 2009; Vogel & Marcotte, 2012). Also, such systems rarely recapitulate endogenous expression patterns and protein function. In this regard, the generation of knock-in lines where the aim is to tag the endogenous locus of a gene with a FP has attracted a lot of attention in the last years. This technique combines the use of genetic tools with the external development of optically translucent embryos making it possible to observe **endogenous** protein expression and dynamics at single cell resolution in real time using live imaging microscopy and thus maximizing the advantages of zebrafish. Furthermore, the use of a knock-in is not limited to the larval stages as it is generally possible to perform immunofluorescence against the FP in dissected tissues and subsequently image.

In zebrafish, only a few KI lines have been reported. The knock-in methods are highly inefficient and vary depending on the region that is targeted, this is the 3’ or the 5’ but also whether the region is an exon, an intron or a non-coding region and whether the reparatory mechanism that is favored after the dsDNA break is homology-directed repair (HDR) or nonhomologous end joining (NHEJ)(Almeida et al., 2021; Hoshijima et al., 2016; Irion et al., 2014; Levic et al., 2021; Wierson et al., 2020). Generally, one could speak of two approaches: 1) protein reporter knock in lines where the endogenous protein is tagged with an FP by creating a fusion protein and 2) transcriptional reporters where a FP is inserted upstream to the ATG disrupting the gene or a FP is connected via a 2A linker to the coding sequence. The last, leads to the generation of two proteins from the same regulatory sequence which can potentially have different turnover characteristics (Gillotay et al., 2020). In this paper we took advantage of a recently published study by the Andersson Group (Jiarui Mi & Olov Andersson, 2023) and developed a *foxo1a* knock in line using CRISPR/Cas9 technology where the coding sequence for EGFP-pA was inserted via homologous recombination in-frame at the 3’ C-terminal of the Foxo1a protein creating a fusion. Briefly, we synthesized 5’ AmC6 modified primers containing a 45bp homology arm to generate a PCR donor that once inserted links the endogenous gene product with a EGFP-pA creating a fusion protein. The donor was coinjected with *in vitro* preassembled Cas9/gRNA ribonucleoprotein complexes (RNPs) targeting the C-terminal to generate mosaic F0 that gave rise to germline transmission (Fig 1A). The selection of our mosaic F0 was based on the fluorescence of the heart under a wide-field fluorescence microscope using GFP channel. However, when we screened those animals in adulthood, they were always giving rise to unfertilized eggs, most probably due to a high number of insertions. Importantly, we also raised F0 injected animals where no apparent GFP+ signal was detected (animals were not evaluated under a confocal microscope) and it was among those animals that we found the F0 with germline transmission. We obtained one germline founder among 30 F0 animals. We validated the precise insertion of EGFP downstream of the coding sequence of *foxo1a* (Fig 1B). We found expression in sprouting as well as developed blood vessels covering the trunk of the animal (Fig 2A & 2B) and the brain (Fig 2C) but not in vessels irrigating deep organs such as the liver (Fig 2D). We also observed expression in the endocardium, epicardium and atrial myocardium (Fig 2E, 2E’ & 2F).The strong expression in the endothelial cells matches well with the fact that in mouse global deletion of FOXO1 is lethal due to incomplete vascular development (Hosaka et al., 2004). At early developmental stages, Foxo1 was also detected in the brain, particularly in the CNS (Fig 2G, 2G’, 2H) as well as in sensory neurons of the olfactory epithelium (Fig 2G, 2G’, 2J), and in the spinal cord in Rohon-Beard (RH) (Fig 2I, 2K’, 2K’’) and Dorsal Root Ganglia (DRG) neurons that arborize below the developing muscles (Fig 2H & 2K). For the brain, studies in mice have shown that *FoxO1* is strongly expressed at mRNA in the striatum and neuronal subsets of the hippocampus (dentate gyrus and the ventral/posterior part of the CA regions) (Hoekman et al., 2006). However, to our knowledge this is the first evidence of expression of *foxo1a* in the developing zebrafish brain. Its characterization requires further work. Furthermore, the retina (Fig 2L & 2L’) and the skeletal muscle (Fig 2A) also express FoxO1.

**Figure 1:**
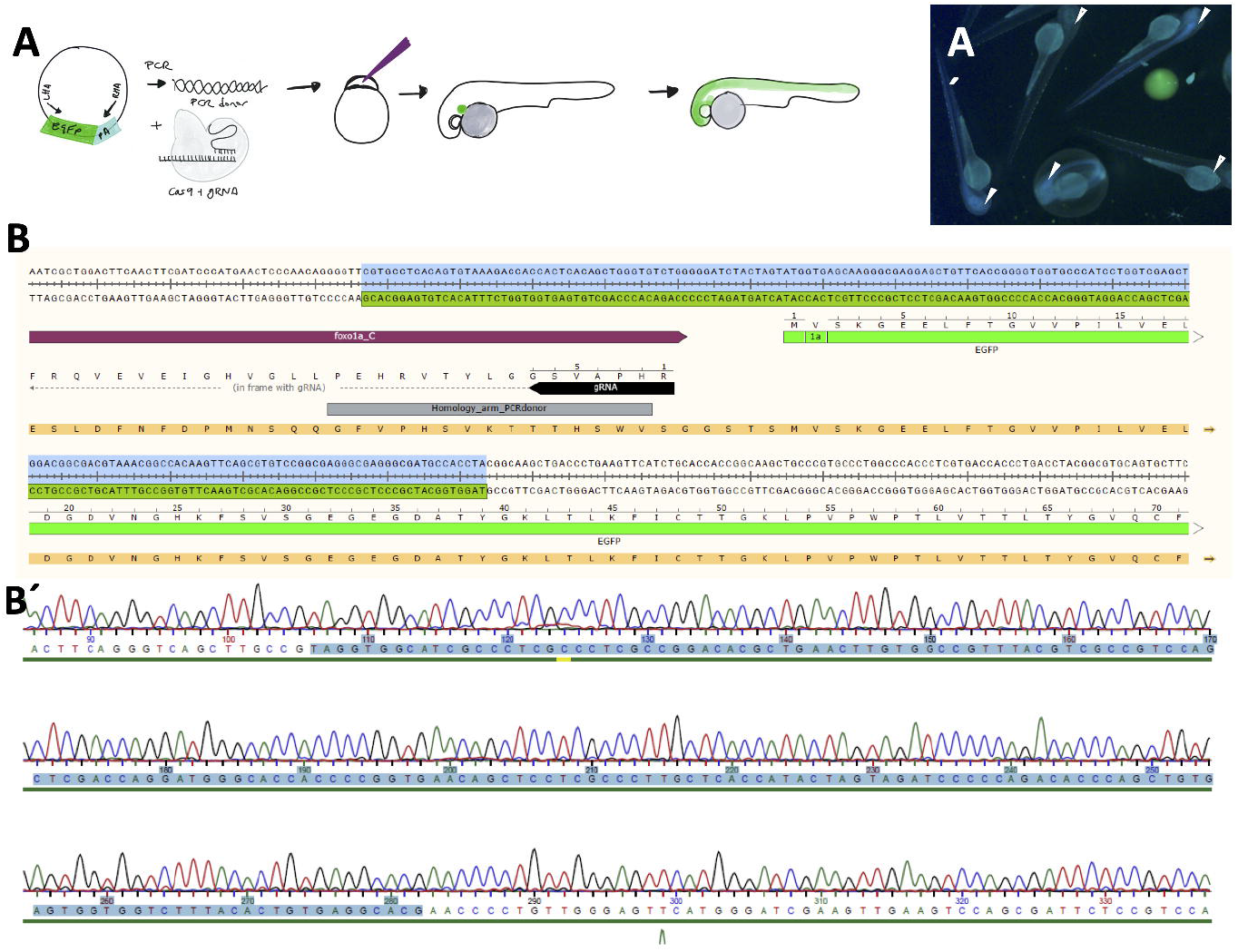
Design and sequencing of the knock in line. (A) Schematic representation of the 3 ’
sknock in strategy. **(A’)** wide-field fluorescence microscope using GFP channel showing F2 positive larvae at 48hpf (white arrows) and negative larvae (empty arrows). **(B)** Target site and insertion of the EGFP-pA cassette at the C-terminal. Note that the protein is a fusion. (B’) Sequencing result verifying the site of insertion (highlighted region in blue).

**Figure 2:**
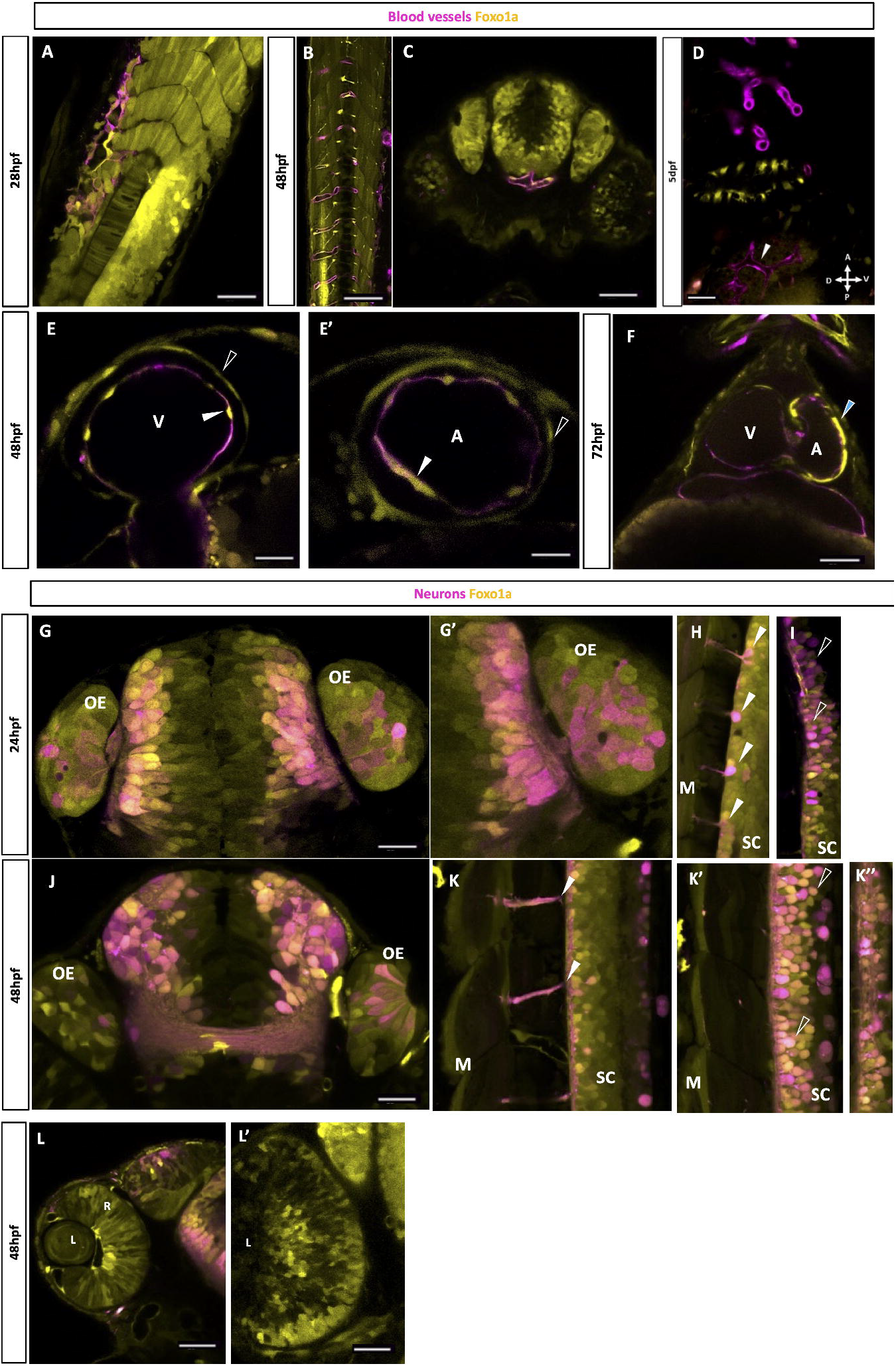
Characterization of the expression of Foxo1a at early developmental stages. (A, B, C, D) Expression of Foxo1a in blood vessels is observed by the co-localization of the Foxo1a EGFP-pA knock in transgene (yellow) and the *Tg(flt1:TdTomato)* line that labels endothelial cells (magenta). **D** shows that endothelial vessels that irrigate the liver do not express Foxo1a (white arrow). Also, note the expression in the skeletal muscle in 2A & 2B. **(E, E’, F)** Expression of Foxo1a in the heart. More specifically in the ventricular and atrial endocardium labelled in magenta (white arrows) and pericardium (black arrows). At 3dpf, the expression is the strongest in the atrial myocardium (blue arrow). **(G, G’, J)**. Co-localization of Foxo1a (yellow) with the transgenic line *Tg(NBT:dsRed)* where the Xenopus neuronal specific beta tubulin promoter drives the expression of DsRed (magenta). Note the expression of Foxo1a in neurons from the CNS but also from the olfactory epithelium (OE). **(H, K)** In the spinal cord (SC) we detect expression in DRG neurons that innervate the muscle (M) (white arrows) and **(I, K’, K’’)** in Rohon-Beard neurons. **(L, L’)** Expression in the retina. All images correspond to in-vivo live imaging under a confocal microscope. Scale bar 50uM.

As a concluding remark, the KI line generated in this paper can be used to characterize the pattern of expression and localization of the protein in a context dependent manner but also opens a big door to explore the FOXO Code, the interacting partners as well as the direct transcriptional targets of this transcription factor. To achieve this, the KI line can be combined with modern techniques like Green CUT&RUN to identify transcriptional targets of Foxo1a (Koidl & Timmers, 2021), with a GFP-directed proteomic mapping combining BioID or TurboID with a GFP-binding nanobody to identify interacting partners (Xiong et al., 2021) and with GFP pull down to perform mass-spectrometry and characterize the FOXO code (Wong et al., 2020). More interestingly, all this can be carried out in different physiological or pathological contexts and either at whole organism level or for a specific tissue or cell type in combination with standard transgenesis and sorting of the cells by flow cytometry.

## MATERIALS AND METHODS

### Zebrafish strains and husbandry

Zebrafish were raised under standard conditions 14 hr light/10 hr dark cycle at 28°C. Adults (90dpf above) were housed in 8L tanks with flow at 28.5°C and embryos up to 5dpf were housed in 8 cm Petri dishes in standard E3 media (5 mM NaCl, 0.17 mM KCl, 0.33 mM CaCl2, and MgSO4) at 28°C (incubated in the dark). All experiments were performed under standard conditions as per the Federation of European Laboratory Animal Science Associations (FELASA) guidelines (Aleström et al., 2020)), and in accordance with institutional Université Libre de Bruxelles (ULB) and national ethical and animal welfare guidelines and regulation, which were approved by the ethical committee for animal welfare (CEBEA) from the Université Libre de Bruxelles (protocols 578N-579N). The following zebrafish strains were used in this study: wild-type (AB), *Tg(−0*.*8flt1:TdTomato)* (G. Wang et al., 2020), *Tg(ins:mCherry-NTR)* (Pisharath et al., 2007) and *Tg(NBT:dsRed)* by Madeleine van Drenth from the Max-Planck Institute for Developmental Biology, Tübingen, Germany. The developmental stages of zebrafish used in experiments are prior to sex specification. All zebrafish were healthy and not involved in previous procedures.

### Generation of the foxo1a-knock in line

For the generation of the knock in line we practically followed the protocol published by Andersson O (ref) which is also discussed here:

#### A. PCR donor design for 3’ knock-in

The dsDNA used for the generation of the KI line also referred to as PCR donor was amplified from a plasmid containing the EGFP-pA sequence using the following primers: The gRNA used is also mentioned.

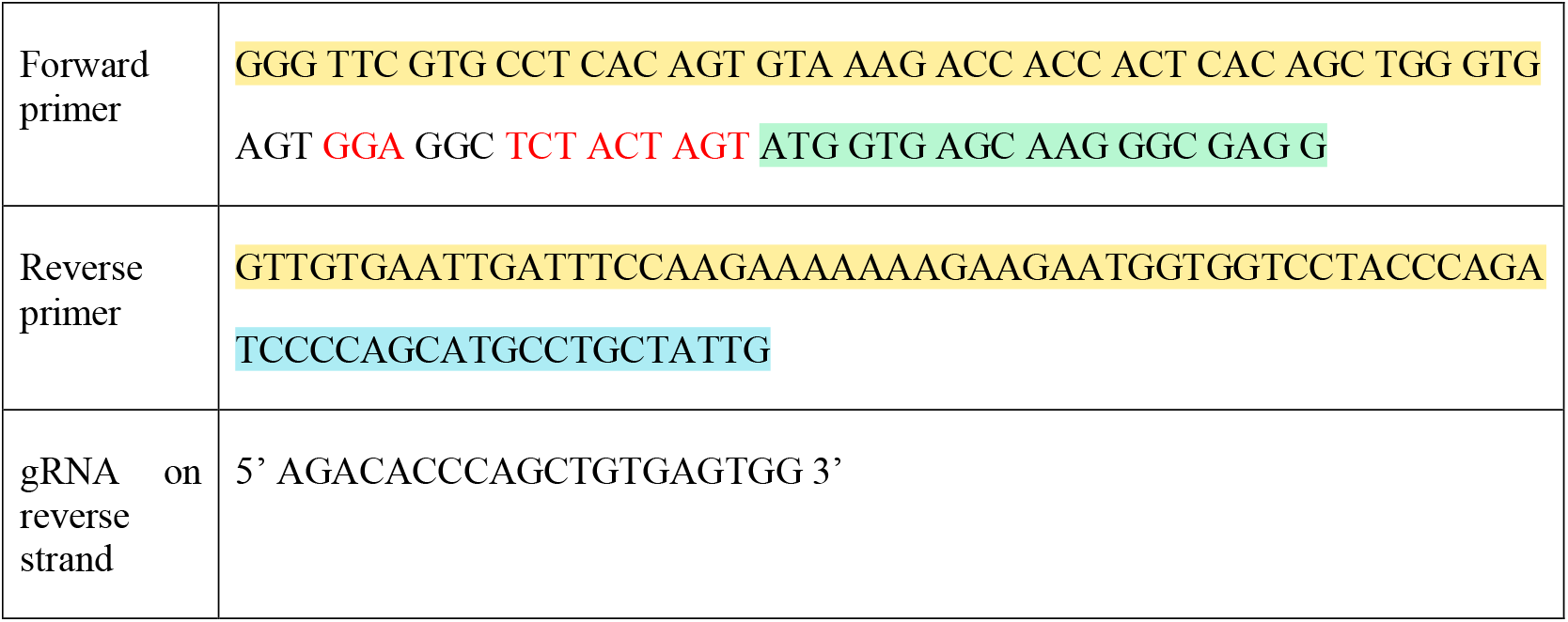

The forward primer includes a homology arm of 45bp (yellow in FP) upstream of the stop codon of the endogenous gene product followed by a linker (only the nucleotides in red are present in the knock-in) and the beginning of the coding sequence for EGFP (green in FP). The reverse primer contains a 45bp homology arm downstream of the endogenous stop codon (yellow in RP) and the end of the bGH polyA sequence (blue in RP).

The 5′ modified primers were ordered with AmC6 5′ modification from Integrated DNA Technologies (IDT). The primer powders were diluted with distilled water into 100 mM as stock solution.

#### B. PCR amplification and gel and column purification

Five PCR reactions of 50 μl PCR mixture were made and amplified using Q5 High-Fidelity DNA polymerase (M0491; NEB). Next, the PCR products were run in 1% agarose gel with 100 V for 45min. The corresponding bands were cut out and pooled all in one tube followed by purification using the Monarch DNA Gel extraction Kit (T1020; NEB) and subsequently repurified using the DNA cleanup kit (T1030; NEB). The concentration of purified PCR product was measured by NanoDrop and stored at -20°C before injection.

#### C. In vitro preassembly of gRNA, Cas9 protein, and donor dsDNA

The Alt-R-modified crRNA, tracrRNA were diluted to 100 μM in low TE buffer, all three ordered from Integrated DNA Technologies and stored at -80°C. A 10uM crRNA:tracrRNA duplex solution was prepared by mixing 10uL of tracrRNA, 10uL of crRNA, 13.3uL of duplex buffer (IDT) and incubating at 95°C for 5min in a thermocycler followed by cooling at room temperature and storage at -80°C. The day prior to the injection the following mix was prepared: 2ul of tracrRNA/crRNA duplex, PCR donor at 20ng/μL, Phenol Red at 0.05%, 1μL of HiFi Cas9 protein (IDT), 5μl of Cas9 working buffer (20mM HEPES, 150mM KCl, pH 7.5) and nuclease free water to a QSP final volume of 20μL. The day of the injection the preassembled mix was incubated at 37°C for 5 minutes and cooled down at room temperature.

#### D. Microinjection and sorting for mosaic F0

We injected the mix into wild-type zebrafish embryos at the early one-cell stage. The overall mortality was around 60% and all dead embryos were removed in the days that followed the injection. At 5dpf, we checked for mosaic F0 based on the fluorescence of the heart under a wide-field fluorescence microscope (LEICA M165 FC) using GFP channel. Positive mosaic F0 were fed rotifers ad libitum from 5dpf to 10dpf and introduced into the fish facility until adulthood. Importantly, we also kept and raised the larvae that did not display any apparent green fluorescence as we hypothesized that animals with low EGFP expression would not be detected in a wide fluorescence microscope.

#### E. Screening and genotyping of F1

At 3 mpf we outcrossed F0 injected animals to wild-type animals. We first started with those that had displayed a mosaic GFP signal in the heart. However, even if these animals were able to lay eggs, all their progeny died within the first 24h most probably due to the high number of insertions. We then started outcrossing animals that had apparently not shown any green fluorescence in F0. After screening around 30 animals, we found 1 animal whose progeny was displaying green fluorescence as early as 24 hpf. At 48 hpf the GFP + progeny was genotyped using a forward primer in the *foxo1a* endogenous locus and a reverse primer in the EGFP inserted transgene using a touch-down PCR with the Terra™ PCR Direct Polymerase Mix (639270; Takara). The PCR product was run in a 1% agarose gel to validate its size and the remaining product was sent for Sanger sequence.

### *In vivo* live imaging

Animals were treated with 5-Propyl-2-thiouracil (PTU; SIGMA) 0.003% in E3 from 24hpf until the day of imaging to prevent pigmentation. PTU was made fresh and renewed every day. The day of imaging, zebrafish larvae were anesthetized in 0.02% of tricaine methanesulfonate (MS-222) (Sigma, E10521) and mounted in 1 % Low-Melt Agarose (LMA) (Lonza, 50080) and imaged on a glass bottom FluoroDish™ (WPI FD3510-100) using a Zeiss LSM 780 confocal microscope. To image the heart, zebrafish larvae were incubated in 0.02% tricaine and 10uM 2,3-Butanedione monoxime (BDM; Thermo Scientific) for 10min prior to mounting in LMA. Also, the agarose cushion was covered with 1mL of 50mM BDM to prevent the heart from beating again while acquiring the images. The structures of interest were imaged using a 40x/1.1N.A. water correction lens. The imaging frame was set at 1024 × 1024 pixels and the distance between confocal planes was set at 6 μm for Z-stack to cover on average a thickness of 120 μm. The samples were excited with 488 nm for Foxo1a, and fluorescence was collected in the range of 487-524 nm and with 543 for dTomato or mCherry signal and fluorescence collected in the range of 570 - 700 nm. Fiji (Schindelin et al., 2012) was utilized for image analysis using maximum-intensity projections of the Z-stack.

## Acknowledgements

We thank the members of IRIBHM fish facility for technical assistance. We thank M. Martens and J.-M. Vanderwinden from the Light Microscopy Facility for technical assistance at ULB.

## Funding

Work was supported by funding from FNRS (MISU-PROL 40005588 and CDR 40013427).

## Author Contribution

Sumeet Pal Singh: conceptualization

Ines Garteizgogeascoa: knock-in generation, confocal imaging, writing.

## Competing interests

The authors declare no competing or financial interests.

## Data Availability Statement

Raw images as well as the knock-in line are available upon request to the corresponding author.

## Notes

### Competing Interest Statement

The authors have declared no competing interest.

## References

Abreu, R. de S., Penalva, L. O., Marcotte, E. M., & Vogel, C. (2009). Global signatures of protein and mRNA expression levels. Molecular BioSystems, 5(12), 1512–1526. https://doi.org/10.1039/b908315d

Aleström, P., D’Angelo, L., Midtlyng, P. J., Schorderet, D. F., Schulte-Merker, S., Sohm, F., & Warner, S. (2020). Zebrafish: Housing and husbandry recommendations. Laboratory Animals, 54(3), 213–224. https://doi.org/10.1177/0023677219869037

Alikhani, M., Maclellan, C. M., Raptis, M., Vora, S., Trackman, P. C., & Graves, D. T. (2007). Advanced glycation end products induce apoptosis in fibroblasts through activation of ROS, MAP kinases, and the FOXO1 transcription factor. American Journal of Physiology. Cell Physiology, 292(2), C850–856. https://doi.org/10.1152/ajpcell.00356.2006

Almeida, M. P., Welker, J. M., Siddiqui, S., Luiken, J., Ekker, S. C., Clark, K. J., Essner, J. J., & McGrail, M. (2021). Endogenous zebrafish proneural Cre drivers generated by CRISPR/Cas9 short homology directed targeted integration. Scientific Reports, 11(1), Article 1. https://doi.org/10.1038/s41598-021-81239-y

Gilels, F., Paquette, S. T., Zhang, J., Rahman, I., & White, P. M. (2013). Mutation of Foxo3 causes adult onset auditory neuropathy and alters cochlear synapse architecture in mice. The Journal of Neuroscience: The Official Journal of the Society for Neuroscience, 33(47), 18409–18424. https://doi.org/10.1523/JNEUROSCI.2529-13.2013

Gillotay, P., Shankar, M., Haerlingen, B., Sema Elif, E., Pozo-Morales, M., Garteizgogeascoa, I., Reinhardt, S., Kränkel, A., Bläsche, J., Petzold, A., Ninov, N., Kesavan, G., Lange, C., Brand, M., Lefort, A., Libert, F., Detours, V., Costagliola, S., & Sumeet Pal S. (2020). Single-cell transcriptome analysis reveals thyrocyte diversity in the zebrafish thyroid gland. EMBO Reports, 21(12), e50612. https://doi.org/10.15252/embr.202050612

Greer, E. L., & Brunet, A. (2005). FOXO transcription factors at the interface between longevity and tumor suppression. Oncogene, 24(50), 7410–7425. https://doi.org/10.1038/sj.onc.1209086

Hoekman, M. F. M., Jacobs, F. M. J., Smidt, M. P., & Burbach, J. P. H. (2006). Spatial and temporal expression of FoxO transcription factors in the developing and adult murine brain. Gene Expression Patterns: GEP, 6(2), 134–140. https://doi.org/10.1016/j.modgep.2005.07.003

Hosaka, T., Biggs, W. H., Tieu, D., Boyer, A. D., Varki, N. M., Cavenee, W. K., & Arden, K. C. (2004). Disruption of forkhead transcription factor (FOXO) family members in mice reveals their functional diversification. Proceedings of the National Academy of Sciences of the United States of America, 101(9), 2975–2980. https://doi.org/10.1073/pnas.0400093101

Hoshijima, K., Jurynec, M. J., & Grunwald, D. J. (2016). Precise Editing of the Zebrafish Genome Made Simple and Efficient. Developmental Cell, 36(6), 654–667. https://doi.org/10.1016/j.devcel.2016.02.015

Irion, U., Krauss, J., & Nüsslein-Volhard, C. (2014). Precise and efficient genome editing in zebrafish using the CRISPR/Cas9 system. Development, 141(24), 4827–4830. https://doi.org/10.1242/dev.115584

Jacobs, F. M. J., van der Heide, L. P., Wijchers, P. J. E. C., Burbach, J. P. H., Hoekman, M. F. M., & Smidt, M. P. (2003). FoxO6, a Novel Member of the FoxO Class of Transcription Factors with Distinct Shuttling Dynamics*. Journal of Biological Chemistry, 278(38), 35959–35967. https://doi.org/10.1074/jbc.M302804200

Jiarui Mi & Olov Andersson. (2023). Efficient knock-in method enabling lineage tracing in zebrafish. Life Science Alliance, 6(5), e202301944. https://doi.org/10.26508/lsa.202301944

Koidl, S., & Timmers, H. T. M. (2021). greenCUT&RUN: Efficient Genomic Profiling of GFP-Tagged Transcription Factors and Chromatin Regulators. Current Protocols, 1(10), e266. https://doi.org/10.1002/cpz1.266

Kops, G. J. P. L., Ruiter, N. D. de De Vries-Smits, A. M. M., Powell, D. R., Bos, J. L., & Burgering, B. M. Th. (1999). Direct control of the Forkhead transcription factor AFX by protein kinase B. Nature, 398(6728), 630–634. https://doi.org/10.1038/19328

Levic, D. S., Yamaguchi, N., Wang, S., Knaut, H., & Bagnat, M. (2021). Knock-in tagging in zebrafish facilitated by insertion into non-coding regions. Development (Cambridge, England), 148(19), dev199994. https://doi.org/10.1242/dev.199994

Pisharath, H., Rhee, J. M., Swanson, M. A., Leach, S. D., & Parsons, M. J. (2007). Targeted ablation of beta cells in the embryonic zebrafish pancreas using E.coli nitroreductase. Mechanisms of Development, 124(3), 218–229. https://doi.org/10.1016/j.mod.2006.11.005

Polter, A., Yang, S., Zmijewska, A. A., van Groen, T., Paik, J.-H., Depinho, R. A., Peng, S. L., Jope, R. S., & Li, X. (2009). Forkhead box, class O transcription factors in brain: Regulation and behavioral manifestation. Biological Psychiatry, 65(2), 150–159. https://doi.org/10.1016/j.biopsych.2008.08.005

Rached, M.-T., Kode, A., Xu, L., Yoshikawa, Y., Paik, J.-H., Depinho, R. A., & Kousteni, S. (2010). FoxO1 is a positive regulator of bone formation by favoring protein synthesis and resistance to oxidative stress in osteoblasts. Cell Metabolism, 11(2), 147–160. https://doi.org/10.1016/j.cmet.2010.01.001

Schindelin, J., Arganda-Carreras, I., Frise, E., Kaynig, V., Longair, M., Pietzsch, T., Preibisch, S., Rueden, C., Saalfeld, S., Schmid, B., Tinevez, J.-Y., White, D. J., Hartenstein, V., Eliceiri, K., Tomancak, P., & Cardona, A. (2012). Fiji—An Open Source platform for biological image analysis. Nature Methods, 9(7), 10.1038/nmeth.2019. https://doi.org/10.1038/nmeth.2019

van der Vos, K. E., & Coffer, P. J. (2011). The extending network of FOXO transcriptional target genes. Antioxidants & Redox Signaling, 14(4), 579–592. https://doi.org/10.1089/ars.2010.3419

Vogel, C., & Marcotte, E. M. (2012). Insights into the regulation of protein abundance from proteomic and transcriptomic analyses. Nature Reviews. Genetics, 13(4), 227–232. https://doi.org/10.1038/nrg3185

Wang, G., Muhl, L., Padberg, Y., Dupont, L., Peterson-Maduro, J., Stehling, M., le Noble, F., Colige, A., Betsholtz, C., Schulte-Merker, S., & van Impel, A. (2020). Specific fibroblast subpopulations and neuronal structures provide local sources of Vegfc-processing components during zebrafish lymphangiogenesis. Nature Communications, 11(1), Article 1. https://doi.org/10.1038/s41467-020-16552-7

Wang, Y., Zhou, Y., & Graves, D. T. (2014). FOXO Transcription Factors: Their Clinical Significance and Regulation. BioMed Research International, 2014, 925350. https://doi.org/10.1155/2014/925350

Wierson, W. A., Welker, J. M., Almeida, M. P., Mann, C. M., Webster, D. A., Torrie, M. E., Weiss, T. J., Kambakam, S., Vollbrecht, M. K., Lan, M., McKeighan, K. C., Levey, J., Ming, Z., Wehmeier, A., Mikelson, C. S., Haltom, J. A., Kwan, K. M., Chien, C.-B., Balciunas, D., … Essner, J. (2020). Efficient targeted integration directed by short homology in zebrafish and mammalian cells. ELife, 9, e53968. https://doi.org/10.7554/eLife.53968

Wong, M., Newton, L. R., Hartmann, J., Hennrich, M. L., Wachsmuth, M., Ronchi, P., Guzmán-Herrera, A., Schwab, Y., Gavin, A.-C., & Gilmour, D. (2020). Dynamic Buffering of Extracellular Chemokine by a Dedicated Scavenger Pathway Enables Robust Adaptation during Directed Tissue Migration. Developmental Cell, 52(4), 492–508.e10. https://doi.org/10.1016/j.devcel.2020.01.013

Xiong, Z., Lo, H. P., McMahon, K.-A., Martel, N., Jones, A., Hill, M. M., Parton, R. G., & Hall, T. E. (2021). In vivo proteomic mapping through GFP-directed proximity-dependent biotin labelling in zebrafish. ELife, 10, e64631. https://doi.org/10.7554/eLife.64631

